# Rates of Karyotypic Evolution in Estrildid Finches Differ Between Island and Continental Clades

**DOI:** 10.1101/013987

**Authors:** Daniel M. Hooper, Trevor D. Price

**Affiliations:** Commitee on Evolutionary Biology, University of Chicago, Chicago, Illinois 60637; Department of Ecology and Evolution, University of Chicago, Chicago, Illinois 60637

**Keywords:** Chromosomes, Estrildid finches, gene flow, inversion, peak shift models, zebra finch

## Abstract

Reasons why chromosomal rearrangements spread to fixation and frequently distinguish related taxa remain poorly understood. We used cytological descriptions of karyotype to identify large pericentric inversions between species of Estrildid finches (family *Estrildidae*) and a time-dated phylogeny to assess the genomic, geographic, and phylogenetic context of karyotype evolution in this group. Inversions between finch species fixed at an average rate of one every 2.26 My. Inversions were twice as likely to fix on the sex chromosomes compared to the autosomes, possibly a result of their repeat density, and inversion fixation rate for all chromosomes scales with range size. Alternative mutagenic input explanations are not supported, as the number of inversions on a chromosome does not correlate with its length or map size. Inversions have fixed 3.3× faster in three continental clades than in two island chain clades, and fixation rate correlates with both range size and the number of sympatric species pairs. These results point to adaptation as the dominant mechanism driving fixation and suggest a role for gene flow in karyotype divergence. A review shows that the rapid karyotype evolution observed in the Estrildid finches appears to be more general across birds, and by implication other understudied taxa.

---

Chromosome inversions are often fixed between closely related species as well as found segregating within species (Coyne and Orr 2004; Hoffmann and Rieseberg 2008; Faria and Navarro 2010). Once established, inversions can foster divergence between populations (Anderson et al. 2005; Lowry and Willis 2010; Jones et al. 2012) and aid in the speciation process (Rieseberg et al. 2001; Noor et al. 2001; Brown et al. 2004; Joron et al. 2011; Ayala et al. 2013; Fishman et al. 2013; Poelstra et al. 2014) by creating barriers to gene flow through the suppression of recombination and/or the induction of structural underdominance in heterokaryotypes. It remains unclear why rearrangements evolve so frequently in some taxa and by what mechanisms, especially as new inversions are often thought to have deleterious fitness effects due to structural underdominance (Stebbins 1958; White 1969; 1978; King 1987; Rieseberg 2001).

Mechanisms proposed to explain the fixation of novel underdominant rearrangements involve both genetic drift and natural selection. As described by Lande (1979), drift within a deme may counter selection and raise the rearrangement above a frequency of 50%, whereupon selection drives it to fixation. Fixation rate in a species is affected by deme size, but not the population size of the species itself. The reason population size does not affect fixation rate is that, after becoming established in one deme, the probability that a rearrangement will fix is equal to the inverse of the total number of demes, but the more demes there are the greater the chance that an inversion becomes established in at least one deme. As in classic models of genetic drift (Kimura 1962) the two effects cancel out and the fixation rate for a species is approximately independent of its population size, although this result depends on assumptions about population structure and the deme where the rearrangement originated (Lande 1979; 1985; Hedrick 1981; Walsh 1982; Spirito 1998). In contrast to the spread of an underdominant rearrangement, if a rearrangement is inherently deleterious its fixation probability within a species should decrease with population size and only realistically occur when population sizes are small (Kimura 1962; Ohta 1972; Whitlock et al. 2004).

Chromosome rearrangements may also become established entirely by positive selection. In this case, a rearrangement can spread to fixation if (1) its breakpoints favorably alter gene expression (Wesley and Eanes 1994), (2) it increases linkage between epistatically interacting sets of genes (Dobzhansky 1951), (3) it increases linkage between locally adapted alleles within a population experiencing gene flow from a divergently adapted population (Charlesworth and Charlesworth 1979; Kirkpatrick and Barton 2006; Feder et al. 2011). In this scenario, gene flow plays a creative role in karyotype evolution because of the selective advantage in keeping locally adapted alleles together. Adaptive models include those external to the individual, as implied in the mechanisms described above, as well as those resulting from genomic conflicts. For example, meiotic drive may favor the spread of a rearrangement if it is associated with a set of interacting alleles that together affect segregation distortion (White 1978; King 1993). Whenever adaptive mechanisms operate, the fixation rate of a novel rearrangement should increase with population size, as the process is mutation-limited. Here, we evaluate fixation rates of large chromosomal inversions in a family of birds, and consider the possible roles of population size and gene flow.

Birds have long been used as models of speciation and are perhaps the best-studied group with respect to how behavior and ecology contribute to population divergence (Price 2008; Grant and Grant 2011) but little attention has been given to the possible role of chromosome rearrangements in bird speciation (Price 2008; Ellegren 2013). This may be because avian genomes are often considered to be highly conserved, and interchromosomal rearrangements such as fusions, fissions, and translocations between species appear to be rare (Takagi and Sasaki 1974; Griffin et al. 2007; Ellegren 2010; 2013; Zhang et al. 2014). For example, diploid chromosome number (2N) for birds typically varies between 76 and 80. However, in some clades 2N is much more variable, notably for finches in the genus *Pytilia* (Christidis 1983), parrots (Nanda et al. 2007), birds of prey (Bed’Hom et al. 2003; de Oliveira et al. 2005; Nanda et al. 2006; Nishida et al. 2008; 2014), and one shorebird, the stone curlew (*Burhinus oedicunemus*; Nie et al. 2009; Hansmann et al. 2009).

In contrast to the general conservation of chromosome number and lack of interchromosomal rearrangements, the cytological literature suggests that intra-chromosomal rearrangements, notably inversions, occur more frequently (reviewed in Shields 1982; Christidis 1990; Price 2008, ch. 16; Völker et al. 2010; Skinner and Griffin 2011; Lithgow et al. 2014, and see Tables 3 and S6 of this paper). These findings are supported by recent comparative genomics studies that report large numbers of inversions (10s to 100s) between distantly related species (Stapley et al. 2008; Hansson et al. 2009; Aslam et al. 2010; Backström et al. 2010; Kawakami et al. 2014; Zhang et al. 2014). After whole genome alignment Kawakami et al. (2014) identified 140 inversions at a median size of ∼2 Mb that distinguish zebra finch (*Taeniopygia guttata*) from collared flycatcher (*Ficedula albicollis*), two species that last shared a common ancestor ∼30 Ma (divergence date inferred from Price et al. 2014). From this, we estimate a fixation rate as fast as one every 0.5 My along the lineage leading to the zebra finch.

Typically a fifth of avian chromosomes are large in size (macro-chromosomes 16060 Mb) and easily distinguishable cytogenetically. The remaining chromosomes are smaller (micro-chromosomes <40 Mb) and the smallest of which are often indistinguishable using standard cytological approaches (Masabanda et al. 2004; Ellegren 2013). Here, we examine the fixation of large cytologically detectable inversions across 32 species of Estrildid finches (order *Passeriformes*). We use published cytological analyses of the Estrildids, which have identified inversion differences between species and polymorphisms within species (Prasad & Patnaik 1977; Christidis 1983; 1986a; 1986b; 1987; Itoh & Arnold 2005). 2N is either 76 or 78 for all Estrildid finches examined except one (the red-winged pytilia, *Pytilia phoenicoptera*, 2N = 56). Seven pairs including the Z can be considered macrochromosomes and the remaining pairs micro-chromosomes. The Z chromosome is the 4^th^ largest while the W is typically 8^th^ in size. The degree of karyotype similarity between species varies considerably and some genera appear to have more labile karyotypes than others (Christidis 1986a; 1986b). A preliminary analysis using a phylogeny for a subset of the karyotyped Estrildid species estimated that the fixation of cytologically detectable chromosomal inversions occurred every 2.7 My along a lineage (Price 2008, pp. 386-388). We expand on this analysis using a time-dated tree to ask how variation in the rate of inversion fixation differs among chromosomes and is associated with demographic and geographic differences between species.

## Methods

The family *Estrildidae* contains approximately 140 species distributed across the Old World tropics and southern hemisphere temperate zone (Goodwin 1982; Sorenson et al. 2004).

The Estrildid finches originated in Africa (Sorenson et al. 2004; Sorenson *pers. comm*.) and their subsequent dispersion and diversification across the Old World have produced 7 discrete radiations that vary extensively in the average population size and degree of sympatry between member species (Goodwin 1982).

## IDENTIFYING INVERSIONS

We use the data of Christidis (1983; 1986a; 1986b; 1987) supplemented by others (Hirschi et al. 1972; Ray-Chaudhuri 1976; Ray-Chaudhuri 1976; Prasad and Patnaik 1977) who describe gross karyotype structure (Tables S1 and S2). These authors used Giemsa staining and C- and G-banding techniques to document large inversions, both pericentric (those encompassing the centromere) and paracentric (not encompassing the centromere). Sample size varied across species (maximum of 20 individuals for *Bathilda ruficauda* and just one for three species) with an average of 5.7 karyotyped individuals per taxon (Table S1).

We converted centromere position and banding pattern for the 6 autosomal macrochromosomes, the 5 largest autosomal micro-chromosomes, and both sex chromosomes into character state data for each species (Table S2). Using published figures, we identified homologous chromosomes between species by chromosome-diagnostic banding patterns and scored them for approximate centromere position (i.e., whether they were metacentric, sub-metacentric, sub-telocentric, or telocentric), following the naming conventions established by Levan et al. (1964). We treated chromosome conformations as distinct when species shared the same general classification (e.g., both were sub-metacentric) but the author of the paper noted that the banding pattern flanking the centromere differed. We identified pericentric inversions between taxa from changes in both the location of the centromere and the orientation of the banding pattern immediately flanking the centromere. We identified paracentric inversions by re-orientation of banding pattern alone. However, as we detected only three paracentric inversions in our dataset and this is likely to be a significant underestimate of their true diversity (Kawakami et al. 2014; Zhang et al. 2014), we therefore only included pericentric inversions in our analyses.

Centromere repositioning can result from processes other than pericentric inversions, such as the redistribution of heterochromatin (Krasikova et al. 2009; Zlotina et al. 2012) and the evolution of neo-centromeres (Marshall et al. 2008). We are able to exclude these alternative explanations because centromere repositioning was matched with inversion of proximal banding patterns.

Five species had polymorphic chromosome conformations due to pericentric inversions found segregating within the individuals karyotyped (Table S2). We did not include them in our analyses because we are only interested in the drivers of inversion fixation. As such, an inversion on a polymorphic chromosome was only counted as a fixed difference between species if the ancestral conformation, determined by Bayesian approach in SIMMAP v1.5 (Bollback 2006) – see below, was not one of the forms found segregating.

## PHYLOGENETIC ANALYSES

Several phylogenetic surveys are available (Sorenson and Payne 2001; Sorenson et al. 2004; Arnaiz-Villena et al. 2009) but there is no complete published phylogeny of the *Estrildidae*. We built a time-dated phylogeny for the 32 karyotyped finches using mitochondrial data from GenBank (Table S3). We dated the tree using the divergence time between the families *Estrildidae* and *Ploceidae* inferred from the multiple fossil-calibrated passerine tree of Price et al. (2014), setting a uniform prior of 17.5-22.1 Ma, based on the 95% confidence limits from the posterior distribution of that tree. Phylogenetic analyses were conducted using BEAST v1.8.0 (Drummond and Rambaut 2007). Data from the mitochondrial loci *cytb* (1143bp), *ND2* (1041bp), and *ND6*+control region (1100bp) were partitioned by locus, each with its own uncorrelated lognormal relaxed clock, and GTR + Γ + I model of sequence evolution (estimated to be the optimal model for each locus using Jmodeltest v0.1.1; Posada 2008). Algorithms were run for 50 million generations and sampled every 5,000 for a total of 10,000 trees of which the first 1,000 were discarded as burn in. We assessed run length and appropriate sampling for each parameter using Tracer v1.5 (Rambaut and Drummond 2007). using TreeAnnotator v1.7.2 (Drummond and Rambaut 2007), we extracted the maximum clade credibility tree, with associated confidence intervals for median node heights.

## INVERSION FIXATION ANALYSES

We estimated the ancestral centromere position (up to 4 possible states: metacentric, sub-metacentric, sub-telocentric, or telocentric) for each chromosome at each node in the tree using the Bayesian approach in SIMMAP v1.5 (Bollback 2006). To account for phylogenetic uncertainty, we simulated over 1,000 randomly drawn trees from the BEAST output post burn in. We obtained the posterior probability estimate for each ancestral centromere position for each chromosome at every node. Inversions were inferred to have occurred upon branches where the karyotype of an internal node differed from subsequent nodes or the tips and was supported by a posterior probability, p > 0.75. The inferred number of inversions per chromosome was concordant with results from reconstructions based on a maximum likelihood model in Mesquite v2.7.5 (Maddison and Maddison 2001; results not shown).

We used ancestral karyotype estimates to calculate the total number of inferred inversions that have occurred on each chromosome separately. Here, we had to exclude three chromosomes from a single species (*Pytilia phoenicoptera*) that have fused and thus precluded identification of chromosome homology (Table S2; excluding this species entirely would not affect conclusions). We next assessed the degree to which the genomic distribution of inversions could be explained by mutagenic input. Under the assumption that the inversion mutation rate is constant per DNA base, the probability of a new inversion on a given chromosome or polytene chromosome arm should be proportional to the chromosome’s physical size. We also assessed the relevance of a chromosome’s map length and GC content based on an alternative process-driven assumption, conditioned on the fact that chromosomal rearrangements are mediated by double-stranded meiotic breaks, that predicts inversions should be proportional to cross-overs (Baudat and de Massy 2007; de Massy 2013). Given that at least one cross-over per chromosome arm appears to be required for proper pairing of homologous chromosomes during meiosis, we assessed whether the fixation rate of inversions was constant per chromosome. Finally, as chromosome breakpoints are often enriched in repeat dense regions (Skinner and Griffin 2011; Kawakami et al. 2014), we assessed whether variation in the number of inversions per chromosome could be explained by variation in repeat content. We estimated chromosome physical size and GC content from the zebra finch genome assembly (Warren et al. 2010), repeat content per chromosome from a RepeatMasker annotation of the zebra finch genome (Smit et al. 1996-2010; http://www.repeatmasker.org), and map distance from two independent zebra finch linkage maps (Stapley et al. 2008; Backström et al. 2010). Hence, we are assuming that the general features of the Estrildid finch genome and recombination landscape (chromosome size, GC content, map distances, and repeat content) are conserved across species with respect to the zebra finch. Comparative studies indicate that chromosome size, GC content, and repeat content varies little even between distantly related avian species (Ellegren 2013; Kawakami et al. 2014; Poelstra et al. 2014; Zhang et al. 2014) and that the recombination landscape has a phylogenetic signal, suggesting our extension of genomic parameters from the zebra finch to this confamilial set of Estrildid finches is warranted (Dumont and Payseur 2008; 2011; Smukowski and Noor 2011). We correlated the number of inversions with chromosome size and map distance, using each chromosome as a replicate. We subsequently compared alternative hypotheses using multiple regression.

In order to examine the influence of demography and geography on the rate of fixation of inversions, we assigned 27 of the 32 Estrildid finches that have cytological data into 5 clades comprising distinct geographic radiations with complete posterior probability support (Figure 1, Table 1, and Figures S1-S5). We focus on clades as they represent distinct monophyletic groupings at a deep timeline. Five species were not assigned to any clade because they were either the only species with cytological data belonging to an independent geographic radiation or were singleton species that did not group with other Estrildid species in their region. For example, the black-and-white mannakin (*Spermestes bicolor*) is the single species with cytological data belonging to a clade that radiated after re-colonizing Africa from Asia, precluding an assessment of inversion evolution within this interesting group. M. Sorenson (*pers. comm*.) provided information on the total number and identity of Estrildid species within each clade (i.e. including species without cytological data), based on unpublished phylogenetic results for the *Estrildidae* (Table 1). The total number of inversions fixed in each clade was summed across chromosomes. We estimated the fixation rate for the clade as the total number of inversions divided by the total branch length connecting all karyotyped species within the clade. We also estimated the pericentric inversion fixation rate, hereafter referred to as the inversion fixation rate, as the total number of pericentric inversions divided by the number of nodes, as a measure of the minimum number of speciation events. This may scale with the number of opportunities for secondary contact between partially reproductively isolated forms between which gene flow may promote the spread of an inversion (Kirkpatrick and Barton 2006).

**Figure 1.**
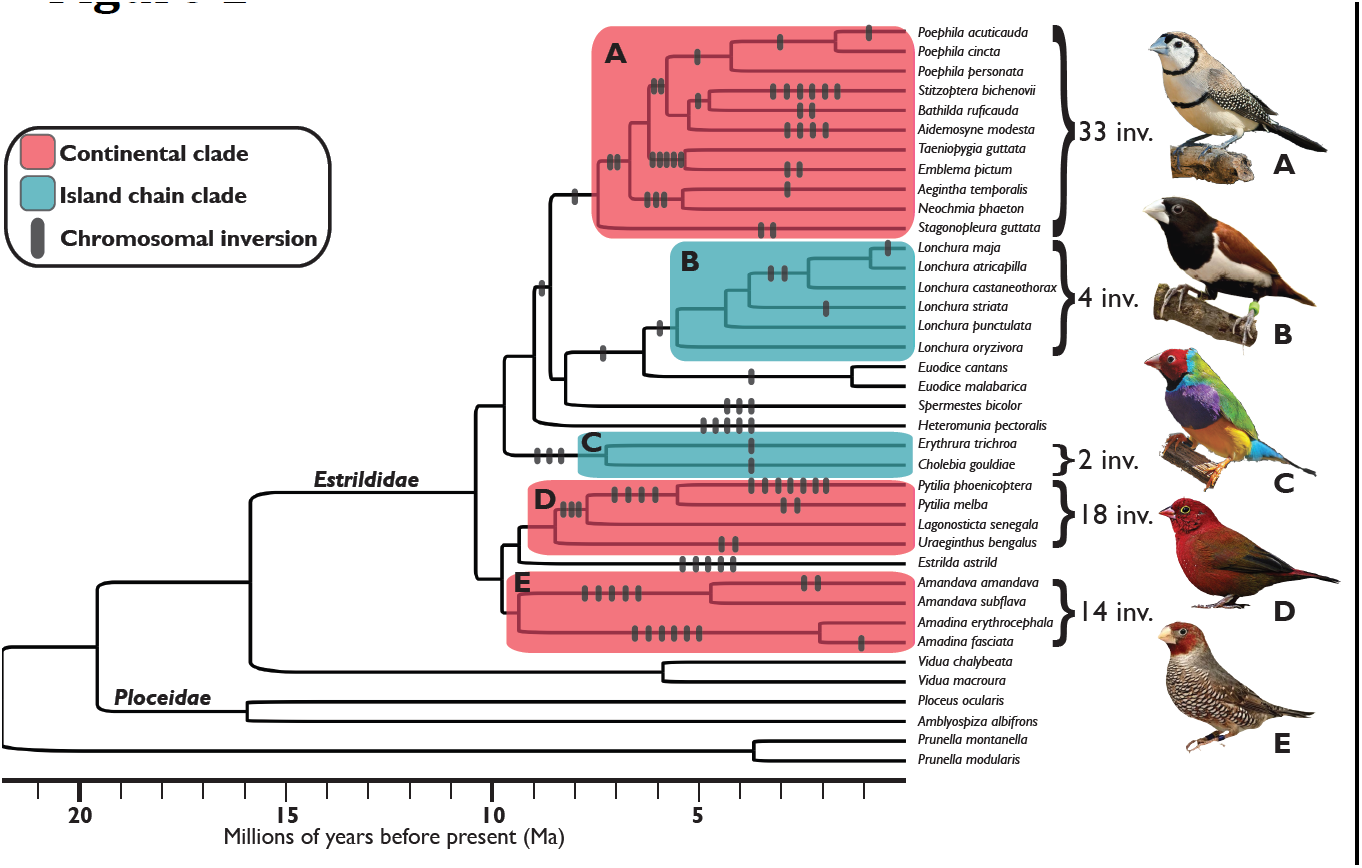
– Phylogenetic distribution of chromosome inversions. Phylogeny of 32 Estrildid finch species with karyotype described and time-dated using the split between families *Estrildidae* and *Ploceidae*. Continental Estrildid clades are boxed in red and island chain clades in blue. The number of inversions inferred to have occurred, based on ancestral karyotype reconstruction, are shown on each branch. The total number of inversions inferred in each of the five clades and a species representative are shown to the right of each clade. All 5 clades have 100% posterior probability support. From top to bottom: (A) grassfinches – *Stitzoptera bichenovii*, (B) munias – *Lonchura malacca*, (C) parrotfinches – *Cholebia gouldiae*, (D) pytilias, cordon-bleus, and firefinches – *Lagonosticta senegala*; (E) adavats, amadina, and quailfinches – *Amadina erythrocephala*.

**Table 1.** – Phylogenetic distribution of chromosome inversions in the *Estrildidae*

| Clade: | Inversion fixation rate (Inv/My) | Number of inversions | Combined branch length (My) | Species karyotyped (Total species) | Crown age (Ma) | Proportion continental species | Avg. Range Size (Range) (10<sup>6</sup> km<sup>2</sup>) | Proportion sympatric* |
| --- | --- | --- | --- | --- | --- | --- | --- | --- |
| **A) Grassfinches** | 0.502 | 31 | 61.72 | 11 (14) | 7.5 | 0.93 | 1.5 (5x10<sup>-2</sup> - 6.3) | 0.57 |
| **B) Munias** | 0.176 | 4 | 22.66 | 6 (26) | 5.6 | 0.23 | 0.99 (10x<sup>-3</sup> - 7.8) | 0.18 |
| **C) Parrotfinches** | 0.137 | 2 | 14.59 | 2 (12) | 7.3 | 0.09 | 0.26 (3x10<sup>-3</sup> - 1.6) | 0.11 |
| **D) Pytilias, Cordon-bleus, & Firefinches** | 0.526 | 16 | 30.4 | 4 (31) | 8.5 | 1.0 | 2.3 (10<sup>-2</sup> - 10.3) | 0.49 |
| **E) Adavats, Amadinas, & Quailfinches** | 0.545 | 14 | 25.68 | 4 (8) | 9.4 | 1.0 | 2.7 (0.5 - 4.0) | 0.39 |

\*Proportion sympatric is the proportion of all pairs of species within each clade that overlapped in range more than 15%.

To assess the relative influences of demography and geography, we extracted species’ range sizes and pairwise range overlaps from natureserve.org using the programs Sp and PBSmapping in R (R Core Team 2014; Tables S4 and S5). We first scored the influence of region (continental vs. island taxa) for each clade as the proportion of species whose ranges were predominately continental. Species were scored as continental or island depending on where the majority of their range lay (Tables 2 and S4). Of the 90 species considered within our 5 clades, only 3 had ranges that were between 25-75% continental (Table S4). Classifying these species alternately as continental or as island did not alter the results in any way.

**Table 2.** – Genomic distribution of pericentric chromosome inversions

| Chromosome ID | Size (Mb) | Map Length (cM)* | GC Content (%) | Repeat Content (%) | Invers |
| --- | --- | --- | --- | --- | --- |
| TGU2 | 156.4 | 34.7 (76) | 39.0 | 0.38 | 3 |
| TGU1 | 118.6 | 63.7 (70) | 39.2 | 0.23 | 8 |
| TGU3 | 112.6 | 22 (69) | 39.4 | 0.38 | 8 |
| TGU1A | 73.7 | 63.3 (91) | 39.7 | 0.14 | 9 |
| TGU4 | 69.8 | 31.9 (18) | 39.2 | 0.28 | 10 |
| TGU5 | 62.4 | 89.8 (64) | 40.8 | 0.12 | 11 |
| TGU7 | 39.8 | 27.8 (41) | 41.1 | 0.14 | 5 |
| TGU6 | 36.3 | 44.6 (60) | 41.6 | 0.19 | 4 |
| TGU8 | 28.0 | 44.9 (47) | 41.3 | 0.05 | 4 |
| TGU9 | 27.2 | 60.5 (52) | 43.1 | 0.15 | 4 |
| TGU12 | 21.6 | 47.5 (34) | 43.7 | 0.07 | 1 |
| Z | 74.6 | 32.8 (29) | 39.2 | 1.5 | 13 |
| W | 27.0 | - | - | - | 11 |

\*Map length values are from the framework map for zebra finch given in Table 2 in St et al. (2008) and those shown in parentheses from the framework map zebra finch m Backström et al. (2010)

Across the *Estrildidae*, range size varies by over 3 orders of magnitude from the red-billed firefinch (*Lagonosticta senegala*), the most widely distributed species (1x10^7^ km^2^), to the red-headed parrotfinch (*Erythrura cyaneovirens*), the most narrowly distributed species (3000 km^2^, Table S4). We use range size as a proxy for population size and assigned a range size score to each clade based on the average range size of all species within it. Range size generally correlates with nucleotide diversity within species, which in turn is correlated with effective population size (Nevo et al. 1984; Cole 2003; Leffler et al. 2012).

In the Estrildid finches, estimates of nucleotide diversity are approximately an order of magnitude greater for zebra finch populations upon the Australian continent compared to zebra finch populations upon the Lesser Sunda Islands, consistent with the two order of magnitude difference in their range size (Balakrishnan and Edwards 2009). It is worth noting that the fixation probability of an adaptive mutation depends on variance in family size (Peischl and Kirkpatrick 2012) and not *N*_*e*_ per se. Variance in family size is only one of several factors that may cause *N*_*e*_ to differ from *N* and we have no reason to suspect that variance in family size varies greatly across Estrildid species.

We define the range overlap of species A with species B as the proportion of species A’s range that is shared with B. Species pairs within clades were scored as sympatric if their average range overlap was greater than 15% (Table S5). Each clade was assigned a sympatry score as the proportion of all possible species pairs that were sympatric. Scoring species’ distributions as sympatric with an average of 10% or 3% range overlap did not qualitatively change the results for the impact of sympatry on inversion fixation rate.

For each factor (region, range size, and range overlap), we calculated the correlation with inversion fixation rate using clade as the replicate. By studying at the clade level (N = 5), we eliminate possibilities for pseudoreplication as far as possible, but the tests are conservative. We fit a multiple linear regression model using fixation rate as the response variable to determine the extent to which variation in karyotype evolution between Estrildid finch clades is affected by demographic and geographic differences. We focus on the rate of inversion fixation (number of inversions / total clade branch length connecting karyotyped species) but also consider number of inversions as a function of the total number of nodes in a clade. Neither of the two alternative methods of defining inversion fixation rate correlates with total branch length (both p > 0.1).

## Results

The time-calibrated phylogeny for those Estrildid finches with cytological data is presented in Figure 1. Our topology matches that of Sorenson et al. (2004) and an unpublished topology of all Estrildid species (Sorenson *pers. comm*.) for all shared nodes. Cytological sampling of each of the 5 fully-supported clades, as determined from a phylogeny of all Estrildids (Sorenson *pers. comm*.), averaged 38% (minimum of 13% in the firefinches, pytilias, and waxbills - clade D - and maximum of 86% in the grassfinches - clade A, Table 1).

The average rate of pericentric inversion fixation across all 32 karyotyped species is one inversion fixed every 2.26 million years (199 My total tree branch length / 88 inferred inversions). The rate varies considerably among clades (Table 1). For example, gross karyotype has changed as fast as one inversion fixed every 175,000 years of evolution (5.72 inversions per million years, Figure 1, clade A) within the Australian grassfinches but has remained identical for over 11 million years of evolution between two species of Indo-Malayan *Lonchura* munias (clade B).

Overall, the rate of inversion fixation differs 13× between the fastest and slowest evolving chromosomes (Table 2). The number of inversions across the 11 autosomes departs from a Poisson distribution (Goodness of fit test, 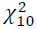 = 93.4) implying that inversion fixation does not occur at equal rates per chromosome. Across the autosomes, the number of inversions per chromosome is not significantly associated with physical size (Pearson’s r = 0.29, p > 0.1), GC content (r = -0.61, p > 0.1), or map length regardless of linkage map used (Stapley et al. 2008: r = 0.38, p > 0.1; Backström et al. 2010: r = 0.3, p > 0.1). Inclusion of the Z further weakened any association.

The two sex chromosomes have fixed inversions at a rate on average twice as fast as the autosomes (two sample t-tests, using the autosomes as replicates: Z chromosome: *t*_*10*_ = 7.1, p < 0.01; W chromosome: *t*_*10*_ = 5.0, p < 0.01). The only significant difference between the autosomes and the Z with regard to the parameters we examined is that the Z chromosome has an order of magnitude greater density of repeats than observed average of the autosomes (two sample *t*-test: *t*_*10*_ = 37.4, p < 0.01; Table 2). There is currently no comparable estimate for repeat content on the W. The W chromosome, however, stands out when comparing chromosomes by the number of inversions per base pair (Table 2). We observe one inversion fixed for every 2.5 Mb on the W, which is over twice as many as on the Z chromosome or the closest autosome TGU5 (both one for every 5.7 Mb).

The fixation rate for inversions is on average 3.3× higher in the three continental than the two island clades (*t*-test: *t*_*3*_ = 16.9, p < 0.01) and is significantly correlated with the proportion of species within clades that have continental distributions (N =5, r = 0.99, p < 0.001). Continental taxa have on average larger ranges and a higher percentage of sympatric species than island taxa (Figures S1-S5). Both average range size (N =5, r = 0.91, one-tailed p = 0.02; Figure 2) and proportion of sympatric pairs (N = 5, r = 0.92, one-tailed p = 0.01) are significantly correlated with inversion fixation rate. In a multiple regression on inversion fixation rate, the partial regression coefficients of range size and proportion of sympatric pairs are both significant (range size: p = 0.042; proportion sympatric: p = 0.037), suggesting that both contribute independently.

**Figure 2.**
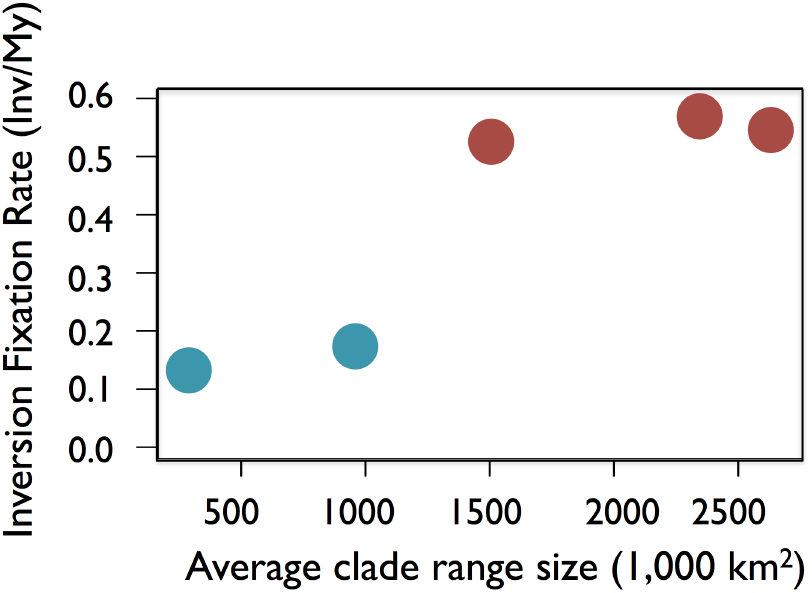
– Rate of inversion fixation (number of inversions / total clade branch length in millions of years) against the average range size of all taxa within each clade. One-tailed Pearson’s correlation: r = 0.91, p = 0.02. Continental Estrildid clades are colored in red and island chain clades in blue. Clades are arranged from left to right (following Table 1): (C) parrotfinches, (B) munia, (A) grassfinches, (D) pytilias, cordon-bleus, and firefinches, (E) adavats, amadina, and quailfinches.

Analyses conducted using inversion fixation rate alternatively defined as the number of inversions within each clade divided by the number of nodes in that clade yielded results similar to those reported above, but they were not as strong and not significant at the P × 0.05 level.

## Discussion

Previous cytological analyses of karyotype structure in Estrildid finches revealed multiple fixed inversion differences between species (Hirschi et al. 1972; Ray-Chaudhuri 1976; Ray-Chaudhuri 1976; Prasad and Patnaik 1977; Christidis 1983; 1986a; 1986b; 1987). We found these inversions have accumulated disproportionately on the sex chromosomes and are much more common within continentally distributed clades, which tend to have larger ranges and a higher proportion of sympatric species. Assuming range size correlates with population size, this is not in accord with the expectations for the fixation of underdominant or deleterious rearrangements because these processes should be independent of (Lande 1979; 1985; Walsh 1982; Spirito 1998) or negatively correlated with (Kimura 1962) a species’ population size. Instead, the correlation with range size suggests that inversions have fixed by positive selection (Whitlock et al. 2004; Vicoso and Charlesworth 2009; Mank et al. 2010b). The positive relationship between range size and inversion fixation rate is present on the autosomes combined (across the 5 clades, r = 0.82) and each of the sex chromosomes (Z: r = 0.44 and W: r = 0.84)

Overall, our results highlight the rapid rate of karyotype evolution in Estrildid finches; with one pericentric inversion fixed every 2.26 My on average. This value is a minimum estimate of the true rate of chromosomal rearrangement. First, many paracentric inversions may have been missed because of the limited number of C- and G- bands per chromosome arm required to infer them and, second, small inversions that cannot be detected cytologically are likely to occur frequently (Kawakami et al. 2014). Paracentric inversions may be less structurally underdominant than pericentric inversions because; in some groups (e.g. *Drosophila*), crossing over within them produces acentric and dicentric recombinants that are removed to the polar bodies during meiosis (i.e. they do not affect female fertility). Whether similar mechanisms to affect the degree of structural underdominance between paracentric and pericentric inversions exist in birds is not known. Small inversions may be subject to similar selective pressures as those examined here but this awaits genomic analysis (e.g. Kunte et al. 2014; Kawakami et al. 2014; Poelstra el al. 2014).

We find that degree of sympatry between species is positively associated with the rate of inversion accumulation, even after range size is accounted for, which may be expected if gene flow between incipient species has contributed to inversion fixation, as in models where rearrangements that capture locally adapted alleles are favored (Kirkpatrick and Barton 2006; Feder et al. 2011). We first consider the genomic distribution of inversions and then the demographic and geographic context of inversion fixation.

We find mixed support for the idea that mutational input has influenced the genomic distribution of chromosome inversions. Chromosome breakpoints are often located within regions that are repeat dense and this appears to be supported by the enrichment of inversions we observe on the sex chromosomes. The repeat content of the Z chromosome is an order of magnitude greater than the autosomal average and has the greatest number of inversions fixed of any chromosome - perhaps suggesting the Z has a greater structural mutation rate. While we do not know the repeat density of the W chromosome, it is believed to be repeat rich due to the reduced power of selection for fixing DNA replication errors (Dalloul et al. 2010; Völker et al. 2010; Kawakami et al. 2014). Alternative mutagenic explanations for the genomic distribution of inversions, however, bear little support. First, chromosome size is not correlated with inversion fixation rate, which is the naïve expectation if structural mutations have a constant rate per base pair. Second, neither a chromosomes map length nor its GC content are correlated with its rate of inversion fixation suggesting an increased number of cross-overs per se does not necessarily result in an increased rate of inversion fixation. While inversion breakpoints may be more prevalent in areas of high cross-over (Skinner and Griffin 2011; Kawakami et al. 2014) and inversions originate from errors during meiotic crossing-over, in the Estrildid finches the fixation rate of inversions cannot be explained by differences between the recombination landscapes of chromosomes alone.

These results contrast with a recent comparative genomic survey of inversion differences between the zebra finch and the collared flycatcher, which found a positive correlation between chromosome size and number of inversions (Kawakami et al. 2014). The difference may reflect the level of resolution possible between studies. Our study only considered pericentric inversions because of the limitations of the cytological methods used to detect them, while the majority of the inversions found by Kawakami et al. (2014) are small (median size of 2.62 Mb and 0.78 Mb in the lineages leading to zebra finch and collared flycatcher, respectively). Smaller rearrangements potentially come with both fewer deleterious effects and smaller selective advantages, resulting in a more even distribution across the genome. Our results are perhaps striking because, in contrast to small inversions, large pericentric inversions seem more likely to carry an intrinsic selective disadvantage due to underdominance and thus suggest that selection was critical in driving their fixation.

A strong result in our dataset is that inversions accumulated twice as rapidly on the sex chromosomes as on the autosomes. While meiotic drive on the sex chromosomes (i.e. sex chromosome drive) is often associated with the establishment of inversion polymorphisms it is unlikely to lead to their fixation because of the deleterious effects of high sex ratio skew (Jaenike 2001). Rather, the scaling of inversion fixation rate on both the Z and W chromosomes with range size suggests positive selection. On the Z chromosome, immediate exposure of recessive mutations to selection and the preferential accumulation of sexually antagonistic genes may make for generally faster rates of adaptive evolution (Charlesworth et al. 1987). This “faster-Z effect” is well documented in birds yet has been attributed to genetic drift rather than selection (Mank and Ellegren 2007; Mank et al. 2010a; Ellegren 2009; Yan et al. 2010; Zhang et al. 2014). However, explanations crediting the predominant role of drift towards the rapid rate of functional divergence on the Z are flawed as they rely on large assumptions regarding the evolution of sex-biased gene expression and avian effective population sizes (Mank and Ellegren 2007; Mank et al. 2010a). As such, the relative contributions of selection and drift towards the faster-Z effect have yet to be properly examined in birds. Regarding the relative rate of inversion fixation, because Z-linked genes diverge in function more rapidly than autosomal genes, at any point in time before reproductive isolation is complete an inversion on the Z chromosome should be more likely to capture two or more alleles locally adapted—either to that population’s habitat or genomic background—than an inversion on an autosome. Thus, inversions on the Z may be more strongly selected for if gene flow is an important mechanism driving their selective advantage.

The W chromosome has more inversions per Mb than any other chromosome we consider and a fixation rate strongly correlated with range size (r = 0.84, p = 0.038). This is surprising as much of the W is devoid of genic content (in the chicken, of the 1,000 active genes on the Z only 10-100 are thought to remain active on the W chromosome, Chen et al. 2012; Ayers et al. 2013), suggesting few adaptive advantages for a recombination modifier. Under positive selection, the fixation rate is ∼2*Nμs*, where *N* is population size, *μ* is the mutation rate, and *s* the selective advantage of the heterozygote (Haldane 1927; Peischl and Kirkpatrick 2012). This formula assumes that variance in family size is Poisson. Among the three parameters (*N*, *μ*, and *s*), *μ* and *s* may help explain the W chromosome’s relatively high fixation rate. We can dismiss an explanation based solely on *N*, as the population size of the W is ¼ that of the autosomes, which should result in a lower fixation rate. While the W chromosome’s mutation rate at the nucleotide level is estimated to be significantly lower than on the autosomes, due to its female-restricted mechanism of germ-line transmission (Ellegren 2013), the structural mutation rate (*μ*) on the W may be greater because of its elevated repeat content (Dalloul et al. 2010; Völker et al. 2010; Kawakami et al. 2014). While the idea that the high number of inversions on the W is due to its repeat density is plausible, chromosome breakpoints are also more often to be found located in areas that have high recombination rates and a high GC content—both of which are lower on the W than the autosomes (Völker et al. 2010; Kawakami et al. 2014; Ellegren 2013; Graves 2014). Finally, selection (*s*) is a composite variable summing across both positive and deleterious effects on fitness. Possibly, *s* is higher on the W chromosome not because inversions have a greater selective advantage than on the autosomes but rather because the costs of structural rearrangements on the W are reduced.

Species ranges are on average 3.6× larger in continental versus island chain clades and this difference is matched by a 3.3× faster rate of inversion fixation. Adaptive models predict this association, but models where drift operates on structurally underdominant deleterious rearrangements do not (Lande 1979; 1985). The usual explanation for the correlation of population size with fixation rate is that the number of rearrangements (or mutations) arising each generation positively correlates with population size (Kimura 1962; Whitlock et al. 2004). However, our analyses across chromosomes based on physical and map size, as described above, argue for an important role for selection beyond raw mutational input. Besides coming into contact more regularly, enabling gene flow that promotes the spread of inversions in adaptive models, species on continents with larger ranges are also more likely to encompass a wider variety of landscapes and climates across which selection might favor alternate allelic combinations between populations connected by gene flow. These are the theoretical starting conditions under which a novel rearrangement that captures sets of locally adapted alleles may spread to fixation (Kirkpatrick and Barton 2006; Feder et al. 2011).

Both the number of inversions and the rate of inversion fixation appear to be correlated with the proportion of sympatric species within Estrildid clades even after accounting for the effects of time and range size. This could be construed as support for inversion fixation as a consequence of gene flow between partially differentiated populations, if extant sympatry between closely related taxa does indeed reflect parapatric divergence. In an extension of the local adaptation model of Kirkpatrick and Barton (2006), chromosome rearrangements are selected to fix between populations upon secondary contact when reproductive isolation is partial but incomplete (Kirkpatrick 2010). Genomescale sequence data from recently diverged taxa suggests that gene flow between incipient species may occur regularly during the speciation process (e.g., Jones et al. 2012; Ayala et al. 2013; Lowery et al. 2013; Eaton and Ree 2013; Cui et al. 2013) but excluding alternative explanations based on incomplete lineage sorting or post-speciation selection is difficult (reviewed in Sousa and Hey 2013; Cruickshank and Hahn 2014).

As in our study, a greater number of chromosome rearrangements have been found in sympatric compared to allopatric taxa in *Drosophila* (Noor et al. 2001), *Helianthus* sunflowers (Rieseberg 2001), *Anopheles* mosquitoes (Ayala and Coluzzi 2005), *Agrodiaetus* butterflies (Kandul et al. 2007), and rodents in the families *Cricetidae* and *Muridae* (Castiglia 2014). One hypothesis, as stated above, is that sympatry reflects historical gene flow between partially reproductively isolated populations that promoted the fixation of these inversions. An alternative is that the greater number of inversion differences observed between sympatric, relative to allopatric, sister taxa occurs because incipient species are more likely to fuse upon secondary contact when they share karyotype structure (Noor et al. 2001; Rieseberg et al. 2001). Previous studies of the phylogeographic context of rearrangement evolution have not evaluated these alternatives (Ayala and Coluzzi 2005; Kandul et al. 2007; Castiglia 2014). If rearrangements accumulate at a constant rate, and incipient species that differ by chromosome rearrangements are far more likely to persist upon secondary contact than those that differ not, then the extent of karyotypic differentiation between sympatric taxa should comprise the upper end of the distribution of karyotypic differentiation between allopatric ones. We do not observe this pattern in the Estrildid finches. Species pairs from island clades, which contain predominately allopatric taxa, have significantly fewer inversions than species pairs from continental clades, which consist of predominately sympatric taxa, regardless of the age of species compared. However, this is confounded with population size. Future examination into the extent of karyotype differentiation between allopatric sister taxa on continents (e.g. the firetails of southwestern and southeastern Australia, gen. *Stagonopleura*), sympatric sister taxa on islands (e.g. the munia species complex of Papua New Guinea, gen. *Lonchura*), and active speciation in hybrid zones should elucidate the extent to which gene flow facilitates inversion fixation more explicitly.

We conclude that karyotype structure across the Estrildid finches is highly variable and appears to evolve rapidly under certain demographic and geographic conditions. Considering that karyotype divergence has been considered unimportant to avian diversification (Ellegren 2010), one question is whether the Estrildid finches are representative of or an exception to the avian rule. In Table 3 we summarized the past 40+ years of cytological research in songbirds, the avian order comprising ∼45% of all bird species, which suggests that chromosome inversions, while variable in number between families, are a pervasive feature of karyotype evolution in songbirds and often involve the sex chromosomes (considering data from 351 taxa across 56 families, see Tables 3 and S6). The upshot appears to be that the *Estrildidae* are not an exception so much in terms of the raw variation in genomic structure between species as they are an exception with respect to the breadth and intensity of taxa so far examined, likely because many Estrildid species are available from aviculture stocks. Our results suggest that alterations of genomic structure may be as important to bird speciation as has been proposed for other better-studied taxa (reviewed in Hoffmann and Rieseberg 2008; Faria and Navarro 2010). That karyotype evolution by chromosome inversion might be a common feature of avian diversification is an exciting prospect for speciation studies and one that should serve to stimulate future research into the extent of genomic rearrangement between species and the evolutionary context in which these changes occur.

**Table 3.** – Summary of karyotypic variation in *Passeriformes* from cytological analyse Songbird families, based on the taxonomy of Jetz (2012), are listed from the top, roug from basal to most derived. Cell entries refer to the number of autosomal chromoson carrying at least one inversion; A refers to an absence of inversions detected. For the chromosomes: P indicates inversions present and A indicates their absence. The *Estr* are in bold. Only families with karyotype descriptions for two or more taxa are show (see Table S6 for a complete dataset listing taxa examined and references).

| Family | Number of<br>taxa karyotyped | Autosomal<br>chromosomes | Chr. Z | Chr. W |
| --- | --- | --- | --- | --- |
| Suborder Tyranni: |  |  |  |  |
| *Tyrannidae* | 13 | 8 | P | P |
| *Furnariidae* | 2 | 2 | A | A |
| *Thamnophilidae* | 4 | 5 | P | A |
| Suborder Passeri: |  |  |  |  |
| Parvorder Corvida: |  |  |  |  |
| *Corvidae* | 16 | 4 | P | P |
| *Laniidae* | 5 | 4 | P | P |
| *Dicruridae* | 2 | A | A | A |
| *Campephagidae* | 2 | 2 | P | P |
| *Oreolidae* | 3 | 1 | P | A |
| *Vireonidae* | 4 | 3 | A | A |
| Parvorder Passerida: |  |  |  |  |
| *Sittidae* | 2 | 2 | P | A |
| *Sturnidae* | 6 | 4 | P | P |
| *Mimidae* | 3 | 2 | P | A |
| *Muscicapidae* | 29 | 7 | P | P |
| *Turdidae* | 16 | 6 | P | P |
| *Bombycillidae* | 2 | 1 | A | A |
| *Chloropsidae* | 2 | A | P | P |
| *Ploceidae* | 3 | 3 | A | A |
| ***Estrildidae*** | **32** | **11** | **P** | **P** |
| *Motacillidae* | 5 | 4 | P | P |
| *Passeridae* | 6 | 4 | P | P |
| *Emberizidae* | 16 | 8 | P | P |
| *Icteridae* | 3 | 3 | P | P |
| *Parulidae* | 6 | 2 | A | A |
| *Passerelidae* | 17 | 8 | P | P |
| *Cardinalidae* | 7 | 2 | P | P |
| *Thraupidae* | 42 | 6 | P | P |
| *Fringillidae* | 21 | 11 | P | P |
| *Paridae* | 5 | A | A | A |
| *Hirundinidae* | 3 | 1 | A | A |
| *Pycnonotidae* | 8 | 6 | P | P |
| *Locustellidae* | 3 | 3 | P | A |
| *Cisticolidae* | 2 | 2 | A | A |
| *Acrocephalidae* | 3 | 5 | P | A |
| *Aegithalidae* | 3 | 2 | P | P |
| *Phylloscopidae* | 9 | 6 | P | P |
| *Sylviidae* | 5 | 5 | P | A |
| *Zosteropidae* | 4 | 1 | A | A |
| *Timaliidae* | 3 | 1 | P | A |
| *Pellorneidae* | 2 | 2 | P | P |
| *Leiothrichidae* | 15 | 6 | A | P |

## ACKNOWLEDGEMENTS

We thank M. Sorenson for sharing some of his data and S. Dubay, M. Kirkpatrick, M. Kronforst, A. Hipp, S. Singhal, Supriya, M. Przeworski, as well as three anonymous reviewers for their advice and assistance with analyses. We declare no conflict of interest for either of the authors in the publication of this paper.

